# Leveraging Segmentation Variability to Improve Brain Age Prediction

**DOI:** 10.64898/2026.07.23.740400

**Authors:** Jacob Sanz-Robinson, Tristan Glatard, Jean-Baptiste Poline

## Abstract

Analytical variability in neuroimaging pipelines contributes to concerns about reproducibility in the field. In structural MRI, different segmentation tools produce discrepant morphometric estimates that may influence downstream analyses, such as predictive modeling. We tested whether integrating several segmentation pipelines improves brain age prediction, and characterized the spatial and demographic structure of pipeline differences across several datasets.

T1-weighted scans from five open-access datasets were processed with four widely-used structural segmentation pipelines. Brain age models were trained using single-pipeline features and compared with multi-pipeline aggregation strategies. Inter-pipeline variability was assessed across shared subcortical structures and examined in relation to age and sex.

Integrating features across distinct segmentation frameworks improved predictive performance relative to individual pipelines, whereas aggregation within closely related software versions provided limited benefit. Variability was spatially structured and volumetric measures were often systematically associated with age and sex.

These results suggest that segmentation differences reflect structured, demographically sensitive variation rather than random noise, and that multi-pipeline feature integration can enhance robustness in neuroimaging-based prediction.

## 1. Introduction

Neuroimaging research has faced persistent concerns regarding reproducibility, with analytical flexibility identified as a major contributing factor (Kennedy et al., 2019). Analytical flexibility refers to the range of outcomes that can result when different, yet common, analysis pipelines, software versions, or parameter settings are applied to the same dataset (Botvinik-Nezer & Wager, 2023). Empirical work has demonstrated substantial variability across analytical choices. For example, thousands of plausible fMRI pipeline configurations can produce divergent activation maps (Carp, 2012), and widely used fMRI software packages can yield limited spatial overlap (Bowring et al., 2019). In the NARPS study, 70 independent teams analyzing the same dataset reached no consensus on five of nine hypotheses, with no two teams using identical workflows (Botvinik-Nezer et al., 2020). Similar multi-team “multiverse” investigations have reported substantial between-group variability in other modalities: in diffusion MRI tractography, 42 research groups segmenting identical white matter pathways produced reconstructions that varied more across analysis protocols than across subjects (Schilling et al., 2021), and in functional Near-Infrared Spectroscopy, 38 independent teams analyzing shared datasets showed low agreement in statistical significance decisions, largely driven by differences in preprocessing and modeling choices (Yücel et al., 2024).

Structural neuroimaging is not exempt from this variability. Widely used segmentation tools such as FreeSurfer (Fischl, 2012) and FSL (Patenaude et al., 2011) can produce systematically different volumetric estimates, with reports of low cross-tool correlations and consistent volumetric offsets (Gomez-Ramirez et al., 2022; Sanz-Robinson et al., 2022). Within-tool variability has also been documented: differences in software version, parcellation scheme, and quality control procedures can lead to statistically distinguishable measures of cortical thickness and regional volume (Bhagwat et al., 2021; Sokołowski et al., 2024).

The choice of cortical parcellation atlas can also influence downstream predictive modeling. Prior studies have shown that prediction performance may vary across parcellation schemes and resolutions (Litwińczuk et al., 2024; Zeighami & Evans, 2021), that atlas selection can affect brain age prediction accuracy (Zhao et al., 2019).

Result variability may further arise from differences in computational environments (Glatard et al., 2015).

Recent work has proposed that analytical variability need not be treated solely as noise. Multiverse frameworks recommend evaluating the stability of findings across plausible analytical choices (Bhagwat et al., 2021; Botvinik-Nezer & Wager, 2023). In the context of structural segmentation, incorporating heterogeneous tool derivatives as features has been shown to improve downstream model robustness. Shuffling outputs across different versions of FreeSurfer improved the performance and generalizability of brain age models, suggesting reduced reliance on version-specific artifacts (Korbmacher et al., 2024). Similarly, numerical perturbation approaches such as Monte Carlo Arithmetic (MCA) have been used to introduce controlled variability into pipelines; training on perturbed outputs improved robustness in brain age and BMI prediction models (Kiar et al., 2021). More broadly, methodological frameworks have argued for leveraging variability to enhance generalization (Kiar et al., 2024).

Brain age prediction provides a suitable setting in which to examine these issues. The brain-age gap—the difference between predicted biological brain age and chronological age—has been proposed as a personalized biomarker of neurodegenerative and psychiatric risk, with elevated gaps reported in clinical populations (Liu et al., 2025; Zhang et al., 2025). Early work applied volume-preserving spatial normalization to structural MRI data and used the resulting regional volumetric maps in Support Vector Machine (SVM) models to classify individuals into age groups (Lao et al., 2004). Subsequent studies shifted toward direct age regression, typically using spatially normalized gray matter (GM), white matter (WM), and cerebrospinal fluid (CSF) maps as features, often combined with dimensionality reduction techniques such as Principal Component Analysis prior to classical regression modeling (Cole et al., 2018; Franke et al., 2010). With advances in segmentation, surface-based morphometry, particularly regional cortical thickness measures, became competitive features for age prediction (Guan et al., 2024; Modabbernia et al., 2022). While early deep learning approaches performed comparably to classical methods (Beheshti et al., 2022), recent large-scale datasets and advanced architectures—including 3D convolutional neural networks (De Bonis et al., 2024) and Transformer-based models (Wu et al., 2024)—have achieved unmatched mean absolute errors (MAEs) in the range of 1–3 years for population age range of 44-85 (Capó et al., 2025; Rajabli et al., 2025).

Despite advances in modeling, feature-based brain age approaches remain dependent on structural segmentation. Systematic differences in volumetric and cortical thickness estimates across pipelines directly alter feature distributions and influence downstream predictions (Bhagwat et al., 2021). Comparisons of commonly used tools have reported substantial disagreement: correlations between cortical thickness estimates from different software packages (e.g., FreeSurfer, CIVET, and ANTs) can be as low as *r* ≈ 0.39–0.52 (Bhagwat et al., 2021), and large systematic offsets have been observed in subcortical volumes (e.g., Cohen’s *d* > 1.4 between FreeSurfer and FSL estimates) (Gomez-Ramirez et al., 2022). Most brain age studies rely on a single segmentation pipeline. This raises a practical question: can integrating outputs from multiple pipelines improve the performance and robustness of brain age models?

At the same time, the structure of segmentation variability itself remains insufficiently characterized. The spatial distribution of inter-pipeline differences across the brain, and their associations with biological variables such as age and sex, are not well understood. Characterizing these patterns provides important context for interpreting both discrepancies between pipelines and the potential benefits of their integration.

In this study, we therefore address two complementary questions. First, do differences between segmentation pipelines capture complementary biological information rather than purely methodological noise? To test this, we evaluate whether integrating multiple pipelines through multi-pipeline aggregation strategies improves the performance and robustness of brain age prediction. We also examine whether variation in cortical parcellation atlas influences model performance by comparing features derived from two commonly used FreeSurfer atlases. Second, how is segmentation variability distributed across the brain, and how does it relate to age and sex?

We processed over 5,000 T1-weighted scans from five open datasets using four widely used structural segmentation pipelines. We quantified spatial overlap and volumetric differences across pipelines and examined their associations with age and sex as descriptive characterizations of variability. We then compared individual pipelines with aggregation strategies, including feature concatenation and regression-based ensemble averaging in brain age prediction tasks.

By systematically characterizing inter-pipeline variability and evaluating its integration in downstream modeling, this work provides practical guidance for improving robustness and reproducibility in neuroimaging-based biomarker development.

## 2. Methods

T1-weighted MRI data from five open-access, lifespan-spanning cohorts were processed using four established segmentation frameworks to extract morphological brain features. These features were used to train and evaluate machine learning models for brain age prediction using nested cross-validation and sex-stratified analyses, including experiments that evaluated the performance of individual pipelines as well as aggregated and ensemble pipeline approaches. Inter-pipeline analytical variability was further quantified using spatial and volumetric agreement metrics across shared subcortical structures, and the influence of demographic factors on volumetric differences was evaluated.

### a) Datasets

To establish a representative benchmark, we utilized five open-access, BIDS-compatible datasets selected for their broad age coverage and multi-scanner variability. This combined sample consists of the community-representative NKI Rockland Sample (Nooner et al., 2012), The PREVENT-AD (Villeneuve et al., 2025) longitudinal cohort of individuals at risk for Alzheimer’s disease, and three healthy volunteer datasets from OpenNeuro: NIMH Intramural Healthy Volunteer (ds005752) (Nugent et al., 2022), Neurocognitive Aging Data (ds003592) (Spreng et al., 2022), and Narratives (ds002345) (Nastase et al., 2021). Figure 1 illustrates the age distribution for all successfully processed sessions (all the pipelines involved completed execution producing the required non-corrupt outputs) with complete demographic data (age and sex), while Table 1 provides comprehensive metadata for each cohort.

**Figure 1:**
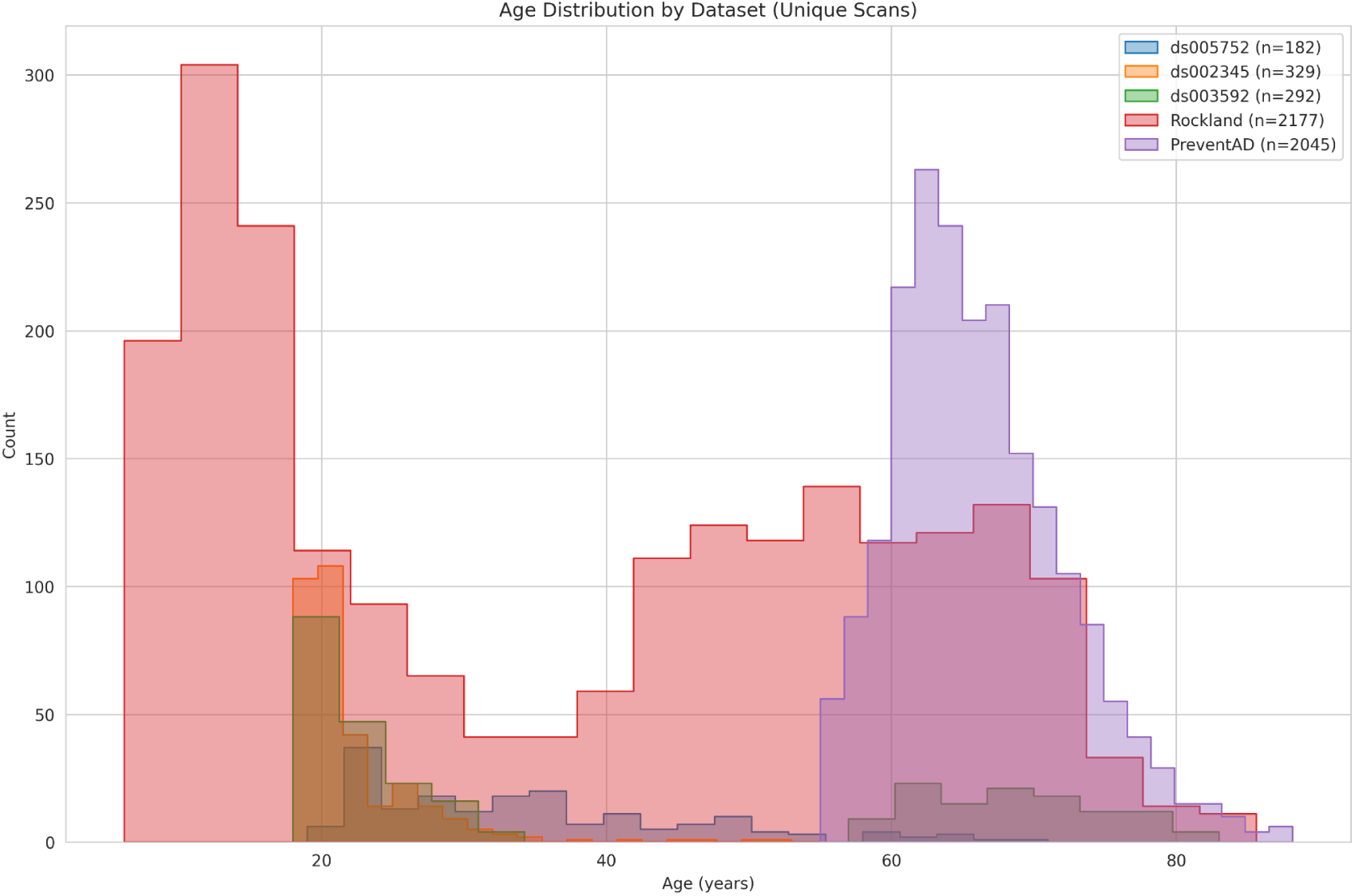
Age distribution for all successfully processed sessions with complete demographic data (age and sex) for the five datasets utilized.

**Table 1:** Summary of the five open-access BIDS datasets used in the study.

| Dataset | No. subjects with T1s | No. T1 Scans | Age range | Description | Scanners |
| --- | --- | --- | --- | --- | --- |
| ds002345 | 345 | 345 | 18-53 | Healthy participants | • Siemens: Magnetom Skyra, Magnetom Prisma |
| ds003592 | 301 | 301 | 18-83 | Healthy participants | • GE: Discovery MR750<br>• Siemens: TimTrio |
| ds005752 | 251 | 252 | 18-71 | Healthy participants | • Siemens: Magnetom |
|  |  |  |  |  | Prisma VE11B-C<br>• Philips: Achieva dStream, Ingenia<br>• GE: MR750, DV25-26 |
| **Prevent-AD** | **348** | **2084** | **55-90** | Longitudinal at-risk Alzheimer's cohort. | • Siemens: TimTrio, Prisma Fit |
| **NKI** | **1330** | **2280** | **6-85** | Community sample of participants across lifespan. | • Siemens: Tim Trio |

### b) Segmentation Pipelines

To evaluate segmentation consistency across different algorithmic frameworks, we utilized four distinct pipelines that are widely-used in the field. All methods are grounded in the Center for Morphometric Analysis (CMA) neuroanatomical segmentation protocol (CMA, 2003; Filipek et al., 1994); however, it is noted that later FreeSurfer versions include specific refinements to individual structures (mainly the thalamic boundaries) (Fogarty, 2018).

The foundational reliance of these four pipelines on the CMA protocol ensures that, in theory, they target an identical set of neuroanatomical structures defined by a unified set of morphological rules. The CMA framework classifies anatomical boundaries into primary intensity-based transitions and secondary knowledge-based subdivisions; the latter requires specific geometric constraints, such as the use of the anterior commissure - posterior commissure line or anatomical landmarks to define ‘cutting’ boundaries where signal contrast is insufficient to separate adjacent gray matter structures. While the core anatomy remains consistent, the practical implications of using different frameworks arise from how each algorithm interprets these primary intensity transitions and operationalizes the secondary manual rules.

The refinements across FreeSurfer—specifically the partitioning of the ‘thalamus proper’ from the ‘ventral diencephalon’—represent a shift in boundary convention rather than a change in the underlying anatomical target. Consequently, systematic variations in segmentations across these four pipelines usually reflect different computational approaches to signal processing and training data within a shared neuroanatomical ontology.

The pipelines utilized were:

- FreeSurfer 7.4.1 (recon-all) (Fischl, 2012): A generative Bayesian model utilizing a Markov Random Field (MRF) prior and a spatially non-stationary Gaussian likelihood term to encode anatomical relationships.
- FreeSurfer 8.0.0.1 (recon-all): A deep-learning framework employing a 3D U-Net trained via Domain Randomization (DR) on synthetic data to enforce the learning of domain-independent features.
- Samseg 8 (Puonti et al., 2016): A parametric Bayesian framework using a deformable tetrahedral mesh and a probabilistic atlas, where intensities are modeled via an unsupervised Gaussian Mixture Model (GMM) that adapts to the input image. Samseg is integrated within the FreeSurfer suite, offering a sequence-adaptive alternative to the traditional recon-all pipeline.
- FSL anat 6.0.7.1 (Patenaude et al., 2011): A Bayesian Appearance Model that estimates the posterior probability of shape given intensity, utilizing learned shape variations as a prior and modeling structures with deformable meshes.

For the FreeSurfer-based pipelines, segmentations were converted from .mgz to NIfTI format using the inbuilt mri_convert tool. Outputs were then registered to 1mm MNI152 space using ANTs 2.4.3 (antsRegistration and antsApplyTransforms) specifically for the subset of downstream tasks requiring a common coordinate system.

### c) Brain Age Regression

We developed two distinct families of models to evaluate the predictive utility of different neuroanatomical feature sets: ‘base’ models and ‘cortical-augmented’ models. We implemented a sparsity filter that excluded any morphological features containing zero or missing values in more than 50% of the samples.

For base models, features were directly derived from segmentations in MNI-152 space across the four segmentation pipelines. While all pipelines provided the 15 common subcortical structures detailed in the subsequent sections, FSL-Anat was limited strictly to these regions. In contrast, FreeSurfer and SAMSEG included a wider array of structures. For these base models, cortical representation was limited to high-level segmented objects, such as the “Left-Cerebral-Cortex.” For each segmented structure, five morphological features were calculated: volume, surface area, compactness, and the first two PCA eigenvalues (length and width). Non-brain matter labels (e.g. Head-ExtraCerebral, WM-hypointensities) were excluded.

For cortical-augmented models, the base models for the FreeSurfer pipelines were augmented with detailed cortical parcellation data. Samseg and FSL anat do not provide these cortical features, so were not used in this family of models. These models incorporated average cortical thicknesses and surface areas derived from the Desikan-Killiany (DKT) and Destrieux (a2009s) atlases.

Predictive modeling was performed using a diverse suite of regressors to ensure a robust evaluation of the data: Elastic Net (EN), K-Nearest Neighbors (KNN), Support Vector Regression (SVR), Extremely Randomized Trees (ET), and Histogram-Gradient Boosting (HGB). A “Dummy” Baseline Predictor was used to provide a performance floor; this model ignores input features and always predicts the mean age of the training set. Crucially, all models were trained and evaluated separately for male and female cohorts to account for sexual dimorphism in brain aging trajectories. This diverse selection of learners was utilized to demonstrate the broad applicability of the multi-pipeline frameworks, rather than to determine the optimal individual predictor.

We employed a Nested 7-fold Group Cross-Validation (CV) strategy. Scans from a single subject remained within the same fold to prevent data leakage. Within each of the 7 outer folds, an inner 3-fold Group CV was utilized for hyperparameter tuning (the specific hyperparameter search spaces for each model are detailed in Supplementary Table S1). Hyperparameter selection was conducted via Randomized Search, evaluating 20 unique parameter combinations per fold.

Data was normalized within the CV loop using a Dataset-Aware Scaler. This calculated the mean and standard deviation for features independently for each dataset/site before scaling, mitigating site-specific batch effects without leaking information from the test set.

Models were evaluated across several configurations (listed in Table 2):

- **Base Pipeline Comparisons:** Individual features from each of the four pipelines separately, a concatenation of all features, and an unweighted ensemble of the four models.
- **Cortical-Augmented Pipeline Experiments:** 1. Single-Atlas: Individual models for each atlas (DKT and a2009s) within each FreeSurfer version. 2. Concatenated Atlases: Merging DKT and a2009s features into a single feature vector for each FreeSurfer version. 3. Four-Way Data Concatenation: Merging the feature sets of all four individual atlas-version combinations (FS7.4.1-DKT, FS7.4.1-a2009s, FS8.0.0.1-DKT, FS8.0.0.1-a2009s) into one high-dimensional dataset. 4. Cortical Ensemble: An unweighted averaging of the predictions from the four individual atlas-version models.

**Table 2:** Summary of all model configurations used in our brain age regression experiments.

| Feature Type | Features per structure | Base Segmentation Pipeline | No. Features |
| --- | --- | --- | --- |
| Base Morphological Features | 5 | FSL | 75 |
|  |  | FS7 | 200 |
|  |  | FS8 | 185 |
|  |  | Samseg | 180 |
|  |  | Data Aggregation | 640 |
|  |  | Ensemble | Multiple |
| Cortical-Augmented Morphological Features | 5 for each base feature, and 2 for each cortical parcellation | FS7-DKT | 324 |
|  |  | FS7-a2009a | 496 |
|  |  | FS7-Both atlases | 620 |
|  |  | FS8-DKT | 309 |
|  |  | FS8-a2009a | 481 |
|  |  | FS8-Both atlases | 605 |
|  |  | Data Aggregation | 1225 |
|  |  | Ensemble | Multiple |

Outlier removal tests were performed using the Median Absolute Deviation (MAD) method (Leys et al., 2013). Unlike the standard deviation, MAD is resilient to extreme values, calculating the median of the absolute deviations from the data’s median. For each feature, a participant was flagged if their value fell outside the range: median ± (threshold x MAD).

Participants were excluded if their total number of flagged features exceeded a specified percentage of the total feature count. We tested two sensitivity thresholds for the base models:

- **Strict:** MAD threshold of 4.0 with a 5% maximum allowed outlier percentage (M=1,325; F=2,555 scans).
- **Moderate:** MAD threshold of 4.0 with a 10% maximum allowed outlier percentage (M=1,511; F=2,830 scans).

The accuracy metric used was Mean Absolute Error (MAE), measured in years. To determine if the top-performing pipeline significantly outperformed the runner-up within the best learner class, we conducted Paired Permutation Testing (N=10,000 permutations). By randomly flipping the signs of these 7 differences, we generated a null distribution of mean differences. This 7-fold resolution provides a discrete p-value floor of approximately p=0.0078, representing the minimum observable significance level for the paired comparison.

### d) Analytical variability analysis: Result discrepancies and relation to subject demography

To quantify inter-pipeline variability, we evaluated 15 common subcortical structures (bilateral Thalamus, Caudate, Putamen, Pallidum, Hippocampus, Amygdala, Accumbens, and the Brainstem) across all four segmentation pipelines. Using a strict session-level intersection, only scans successfully processed by every pipeline were included to maintain a consistent sample size within each figure. Here, successful processing is defined as the completed execution of the pipeline producing the required non-corrupt outputs. No additional quality control was applied to exclude scans, as the aim was to characterize variability among pipelines under typical processing conditions; imperfect segmentations that do not trigger pipeline failure remain part of the variability encountered in practice, and removing them could introduce pipeline-specific selection biases.

Spatial agreement was assessed through voxelwise disagreement heatmaps and Dice-Sørensen Coefficient (DSC). To evaluate the robustness of our DSC overlaps, we performed validations to determine if they were influenced by anatomical scale or specific software architectures, using Bonferroni-corrected Spearman correlations to assess volumetric bias. For the disagreement heatmaps, MNI-space segmentations were masked to include only the subcortical structures of interest. Voxelwise disagreement was computed for each pipeline pair using a multi-class inequality check; a voxel was marked as “disagreeing” if the two pipelines assigned it different structural labels or if only one pipeline assigned a subcortical label. These individual disagreement maps were then averaged across all subjects to yield a global map representing, at each voxel, the proportion of the cohort for which the pipelines disagreed on anatomical identity.

Volumetric consistency was quantified using Spearman rank correlations and Relative Volume Differences (RVD), defined as the absolute difference between pipeline volumes divided by their mean. This metric provides a standardized unit that normalizes for head size and absolute structure volume. Baseline inter-pipeline bias was established via cohort mean RVDs per structure-pair, supplemented by histograms of the volumetric distributions.

Pairwise inter-pipeline differences in volumetric distributions were assessed for each structure using two-sample Kolmogorov–Smirnov tests. Distributional overlap quantified as 1−D, where D is the empirical test statistic. To account for the paired nature of the data (where multiple pipelines processed the same subjects), systematic differences in central tendency were evaluated using two-sided Wilcoxon signed-rank tests, and Bonferroni correction was applied across all structure-wise pipeline comparisons.

Finally, the influence of biological covariates on RVD was evaluated using Spearman’s rank correlation (age) and independent samples t-tests with Cohen’s d (sex). P-values were Bonferroni-corrected across the pipeline pair-structure combinations, with significant findings (ɑ < 0.05) displayed in figures. We performed validations for RVD using Bonferroni-corrected Spearman correlations to assess whether disagreement varied systematically with anatomical size.

### e) Computation Workflow

Orchestrating four segmentation pipelines across five large neuroimaging datasets required coordinated execution across heterogeneous HPC environments, standardized data organization, and systematic tracking of pipeline outputs and downstream analyses. This workflow was managed using NeuroCI v2.0 (Sanz-Robinson et al., 2022), an HPC-oriented orchestration framework designed to automate the execution and comparison of multiple neuroimaging pipelines across datasets. NeuroCI requires BIDS-formatted data and operates on datasets structured according to the Nipoppy (v0.4.0) schema (Bhagwat et al., 2026), which provides the standardized dataset layout and derivative structure on which NeuroCI performs pipeline orchestration and task automation.

The Prevent-AD dataset was processed on the McGill McConnell Brain Imaging Centre (BIC) HPC using NeuroCI, while the remaining four datasets (NKI, ds002345, ds003592, ds005752) were executed directly via Nipoppy on the AllianceCan Rorqual HPC. These Nipoppy-formatted datasets, including all derivatives, were subsequently transferred to BIC and imported into the NeuroCI instance for volumetric and demographic analysis, establishing a reproducible audit trail of artifacts on GitHub. More intensive computations, including segmentation-based disagreement analysis and DSC computations were performed on a local workstation (Intel i7-8650U @ 1.90GHz, 8 cores, 24GB RAM). Regarding machine learning, base morphological models and permutation tests were executed locally, whereas cortical-augmented models were run on the Rorqual HPC. All code is available at https://github.com/neurodatascience/NeuroCI and https://github.com/neurodatascience/pipeline-variability-analysis-scripts.

## 3. Results

### a) Analytical Variability Analysis

#### i) Boundary Delineation Drove Inter-Pipeline Variability and Spatial Overlap Differences

To characterize the spatial variability of subcortical segmentations, we assessed the anatomical distribution of inter-pipeline label discrepancies. Voxelwise analysis revealed that disagreement was consistently low (mean: 0.003–0.005) and primarily localized at structural interfaces and outer boundaries, rather than within the core volumes of subcortical structures (see Supplementary Figure S2 for detailed heatmaps).

To quantify spatial segmentation consistency across pipelines, mean DSCs for the 15 subcortical structures are presented in Figure 2. Spatial overlap varied by structure and pipeline pairing, ranging from 0.55 (FreeSurfer 7 vs. FSL on the left accumbens) to 0.92 (multiple pairs on the brainstem), and shows approximate hemispheric symmetry.

**Figure 2:**
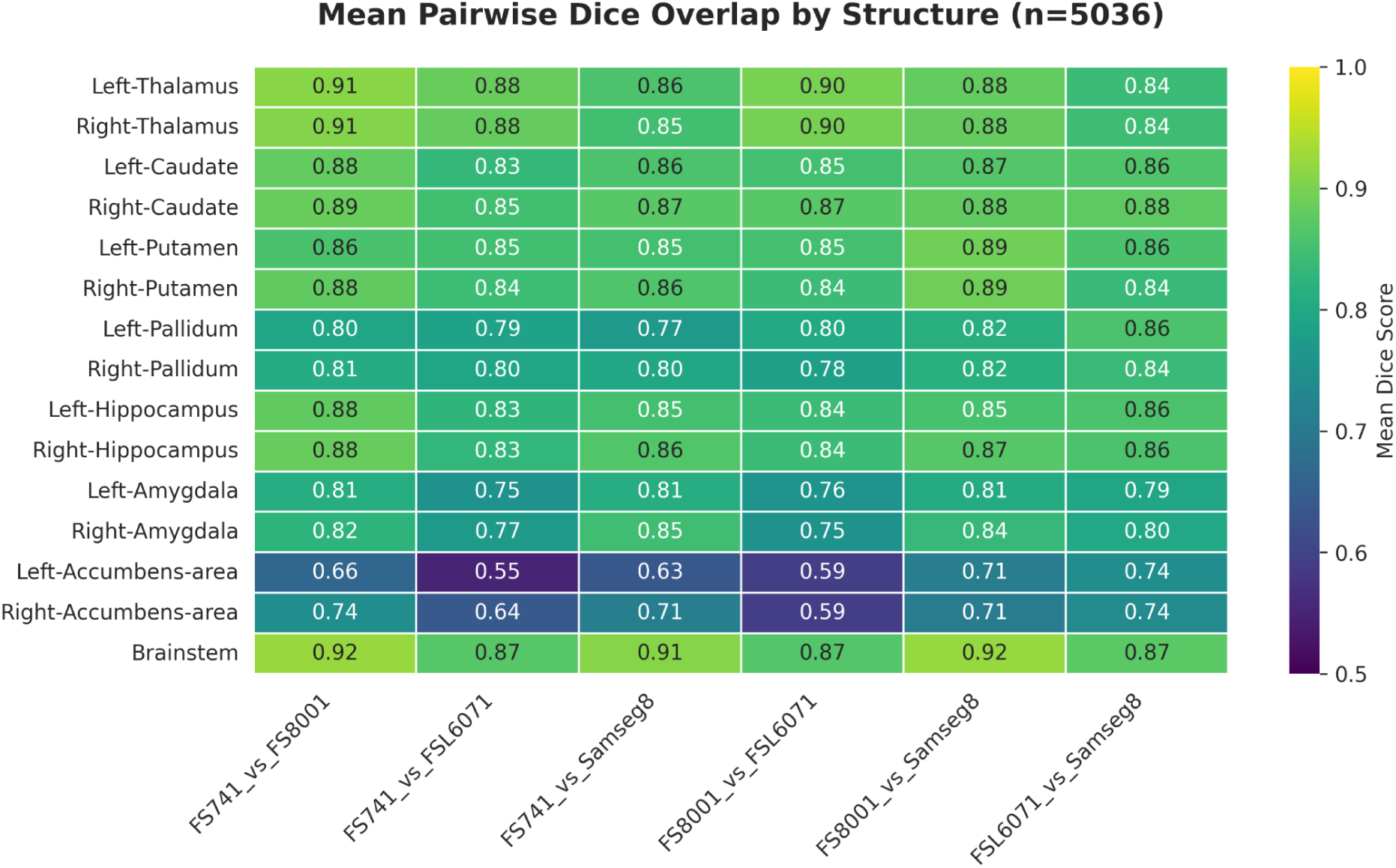
A heatmap showing Dice-Sørensen overlap scores computed across the pipeline pairs for 15 subcortical structures.

As expected given the size dependence of the Dice similarity coefficient, the relationship between structure volume and DSCs was consistently positive across pipeline combinations (Spearman rho range: 0.47–0.93; mean rho = 0.81). After Bonferroni correction for six comparisons, differences within five of six pipeline pairs remained statistically significant. The only exception was the FSL–SAMSEG pairing (rho = 0.47; adjusted p = 0.441), which showed a moderate but non-significant association.

#### ii) Volumetric inter-pipeline disagreement significantly correlated with age and sex

To assess the consistency of volume estimates, we first performed pairwise Spearman rank correlation analysis between pipeline pairs for each subcortical structure, which revealed a moderate-to-high overall agreement across regions (mean ρ = 0.7821, range [0.32, 0.97]; see Figure 3). The highest individual correlation was observed between FS8001 and SAMSEG8 in the brainstem (ρ = 0.9702), while the lowest was found between FSL6071 and FS741 in the left amygdala (ρ = 0.3187).

**Figure 3:**
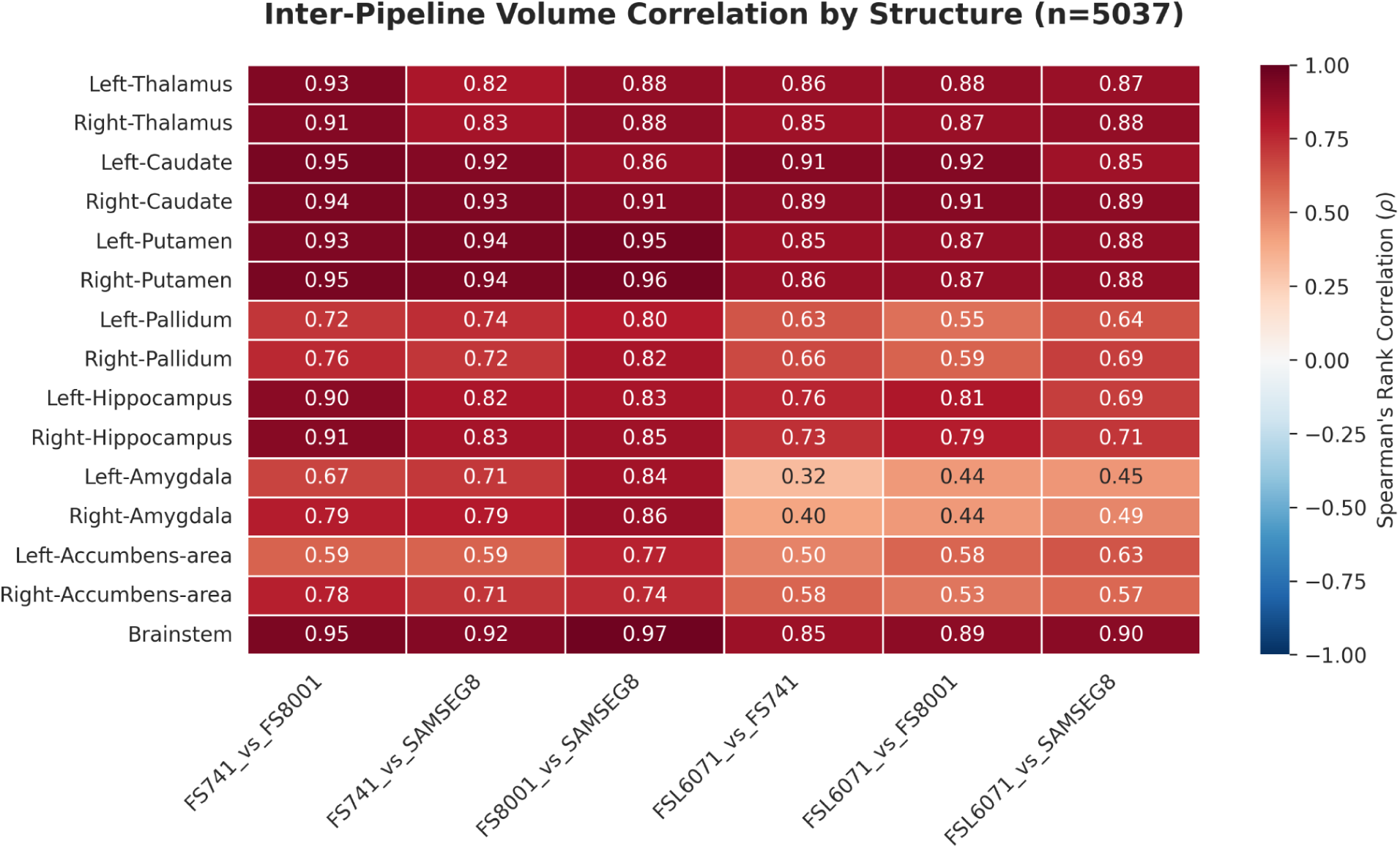
A heatmap showing Spearman rank correlations computed across the pipeline pairs for 15 subcortical structures.

We then compared the volumetric distributions of the 15 subcortical structures across pipelines (see Supplementary Figure S3). Two-sample Kolmogorov–Smirnov tests revealed that no structure exhibited complete distributional equivalence, with mean distributional overlap (1−D) ranging from 0.558 to 0.882. While larger structures like the caudate generally showed higher overlap, the thalamus exhibited only moderate agreement (0.592–0.601) despite its size. Furthermore, the two-sided Wilcoxon signed-rank tests indicated significant systematic differences in central tendency for the vast majority of pipeline pairings.

To further quantify these discrepancies, we computed pairwise RVDs for each structure (Figure 4A). Relative differences were defined as the absolute inter-pipeline volume difference normalized by the mean structure volume, enabling comparison across structures of different sizes. Mean RVDs varied substantially across structures, ranging from approximately 0.04 to 0.42. The accumbens demonstrated high mean RVD (∼40%) across multiple pipeline pairs.

**Figure 4:**
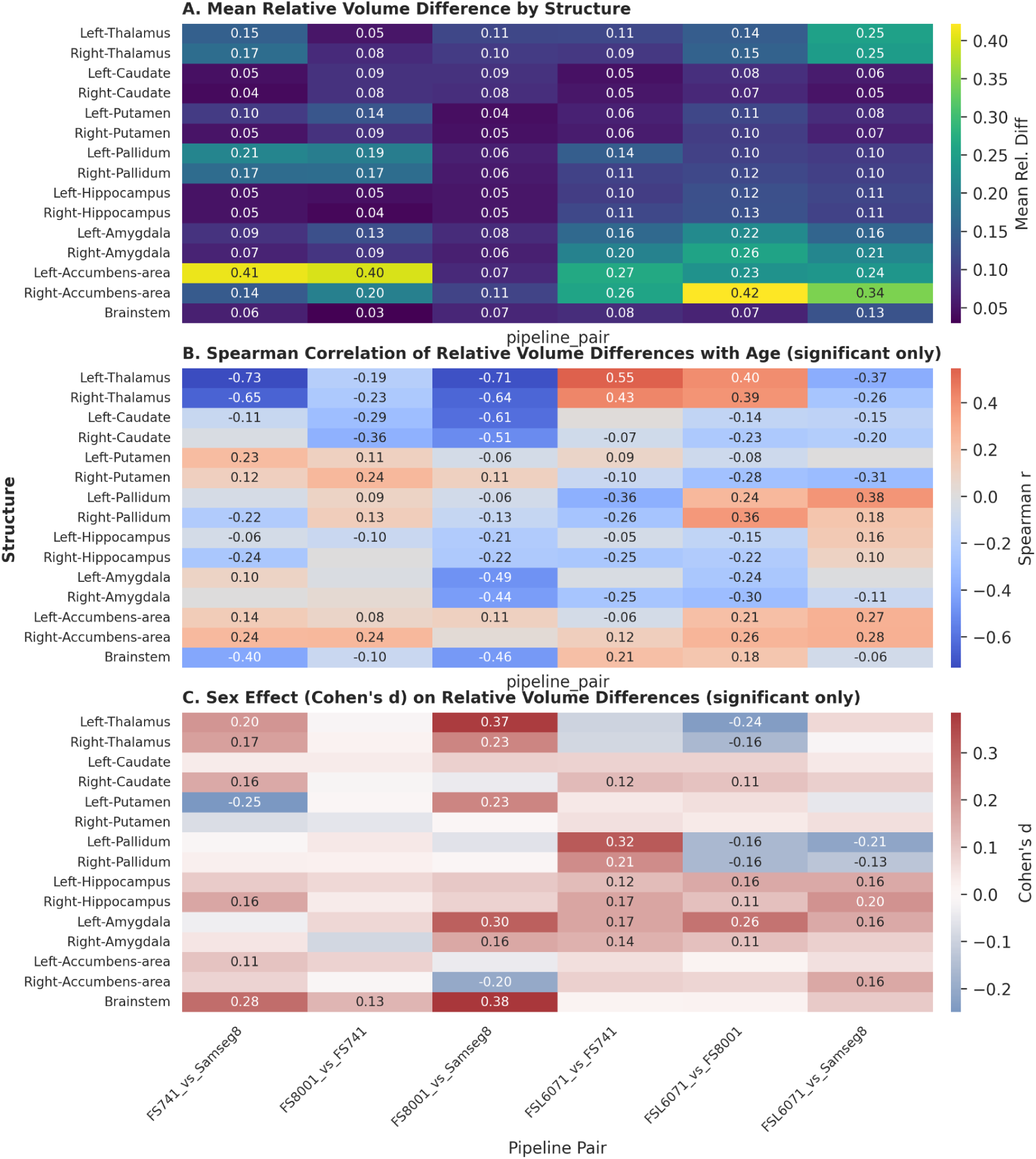
Three separate heatmaps across pipeline pairs for 15 subcortical structures showing pairwise relative volume differences (A), their association with age (B), and sex effects (C).

Spearman correlations between structure-wise median volume and median RVD across six pipeline pairs ranged from −0.732 to 0.018 (mean rho = −0.351). Five of six correlations were negative; one was near zero and positive (FreeSurfer8001-Samseg8001). After Bonferroni correction, only FreeSurfer741-FreeSurfer8001 remained significant (rho = −0.732; adjusted p = 0.011). Overall, evidence for size-dependent disagreement was limited.

Associations between inter-pipeline disagreement and age are shown in Figure 4B. Spearman correlations revealed numerous significant structure- and pipeline-specific effects following multiple-comparison correction, spanning r = −0.73 to r = 0.55. Both the strongest negative and strongest positive correlations were observed in the thalamus across different pipeline comparisons. The direction and magnitude of age associations varied by structure and pipeline pairing. While most structures demonstrated approximate hemispheric symmetry in age-related effects, exceptions were observed in the accumbens and putamen.

Sex-related differences in inter-pipeline disagreement are presented in Figure 4C. Welch’s t-tests were used to accommodate unequal variances, and effects are summarized using Cohen’s d for interpretability. Effect sizes ranged from d = −0.25 to d = 0.38, where positive values indicate greater disagreement in males and negative values indicate greater disagreement in females. Larger disagreement was more commonly observed in males. Significant effects demonstrated consistent hemispheric symmetry. Because Cohen’s d assumes equal variances, these values should be interpreted as approximate indicators of direction and relative magnitude.

Overall, volumetric inter-pipeline disagreement varied substantially by structure and pipeline pairing and showed many significant associations with both age and sex.

### b) Brain Age Regression

#### i) Feature concatenation across pipelines improved base model accuracy

Figure 5 presents the brain age regression results for the ‘base’ models trained on morphological features extracted directly from segmentation outputs (NIfTI files). The data aggregation approach was generally advantageous, with concatenation of features across different pipeline outputs significantly outperforming single-pipeline models in almost all cases, with the exception of the SVM learner for males where slightly better results were obtained with FreeSurfer 8.0.0.1. Applying the same analyses after removing outliers produced nearly identical results, indicating that the observed performance patterns were not driven by extreme cases.

**Figure 5:**
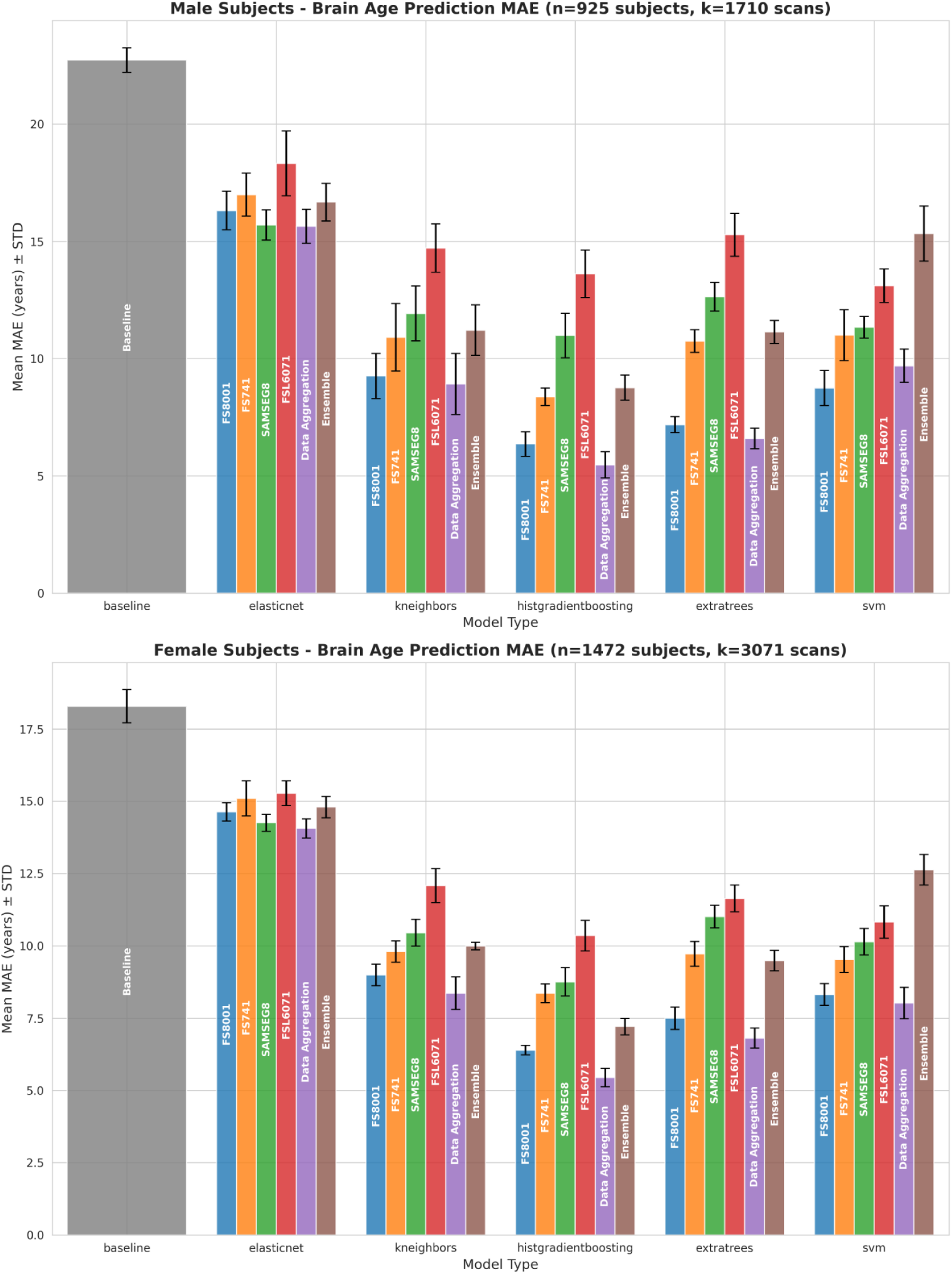
Brain age regression results for our base morphometric models.

The HistGradientBoosting (HGB) model utilizing feature concatenation (Data Aggregation; 640 features) achieved the highest accuracy overall, with a Mean Absolute Error (MAE) of 5.45 ± 0.32 years for females and 5.47 ± 0.56 years for males. This represents a substantial improvement over the baseline (females: 18.28 MAE; males: 22.73 MAE). Although the improvement of the female aggregated HGB model over the best single-source pipeline HGB model (FreeSurfer 8.0.0.1; MAE = 6.39) was modest in absolute terms, it was statistically significant according to paired permutation tests (p = 0.017).

Across all configurations, tree-based architectures (HGB and ExtraTrees) consistently outperformed the linear ElasticNet model. In the context of data aggregation, tree-based architectures also consistently exceeded the performance of distance-based (KNN) and kernel-based (SVM) models. The ensemble method demonstrated comparatively poor performance across learners.

Across all models, prediction accuracy was higher for females (average MAE ≈ 11.5 years) than for males (≈ 13.3 years), likely reflecting both the larger female sample size (N = 3,071 vs. N = 1,710) and the lower age variability in the female group (age SD ≈ 21.0 years vs. ≈ 24.1 years for males), consistent with the lower MAE observed in the baseline dummy model.

Superior performance among single-pipeline models was not solely driven by feature dimensionality. FreeSurfer 8 (185 features) outperformed FreeSurfer 7.4.1 (200 features) across all five learners and both sexes, with the most substantial improvements observed in HGB (MAE = 6.36 vs. 8.37 for males) and ExtraTrees (MAE = 7.18 vs. 10.75 for males). Feature efficiency was further evident in ElasticNet models, where SAMSEG (180 features) achieved lower error than FreeSurfer 7.4.1 (200 features) for both males (MAE = 15.70 vs. 16.99) and females (MAE = 14.25 vs. 15.10). Notably, for male SVM models, FreeSurfer 8 yielded superior accuracy (MAE = 8.75) compared to the higher-dimensional aggregated pipeline (640 features; MAE = 9.69), suggesting that the inclusion of additional features may introduce noise for certain learners.

Feature concatenation, using the largest feature set (n = 640), generally enhanced robustness across folds relative to ensemble methods and lower-dimensional individual pipelines. In male models, concatenation (average SD = 0.75) was more stable than the stacking ensemble (SD = 0.81) and individual pipelines including SAMSEG (n = 180, SD = 0.77), FreeSurfer 7.4.1 (n = 200, SD = 0.86), and FSL-Anat (n = 75, SD = 1.01). Similarly, in female models, the concatenated set (n = 640, SD = 0.42) improved stability over FreeSurfer 7.4.1 (SD = 0.44) and FSL-Anat (SD = 0.51), while matching SAMSEG (SD = 0.42).

Overall, while performance gains over the best single pipeline models were often incremental, the results support the utility of data aggregation as a reliable strategy for refining brain age estimation when using base morphological features, and therefore the hypothesis that different pipelines capture different anatomical signals.

#### ii) Pipeline version and atlas feature concatenation did not improve cortical-augmented model accuracy

Figure 6 presents results for the cortical-augmented models, which incorporated both atlas aggregation (concatenating DKT and a2009s parcellations) and pipeline version aggregation (FreeSurfer 7.4.1 and 8.0.0.1). Across all learners, neither within-version atlas concatenation nor cross-version pipeline aggregation consistently outperformed single-pipeline, single-atlas models. No single pipeline version or atlas demonstrated consistent superiority in either accuracy or fold stability.

**Figure 6:**
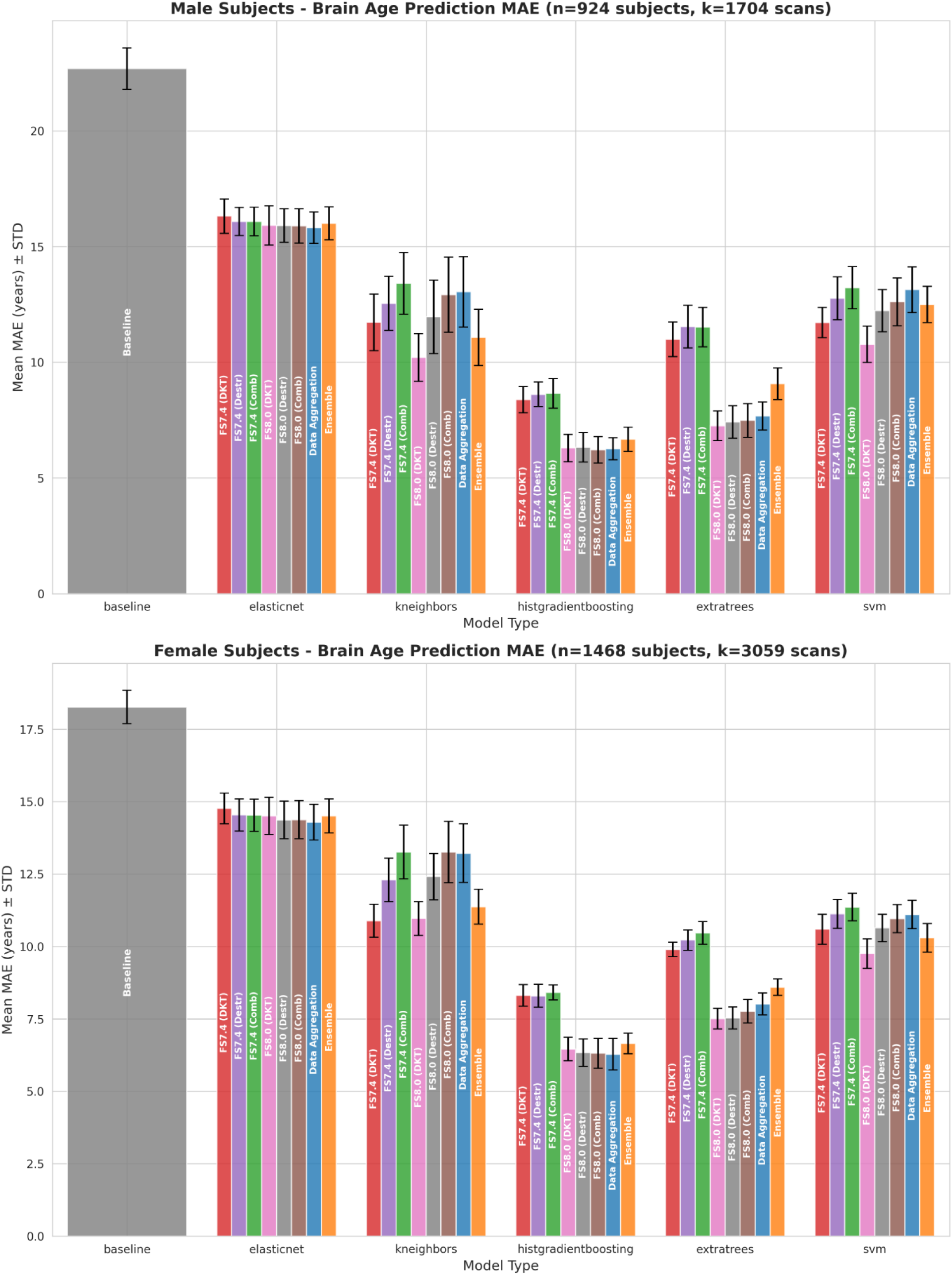
Brain age regression results for our cortical-augmented models.

Tree-based models again provided the strongest performance, and predictions remained more accurate for females (average MAE ≈ 11.80 years) than for males (≈ 13.05 years). HGB was the best overall learner for both sexes. The highest-performing configuration was the HGB model using the FreeSurfer 8.0.0.1 combined atlas configuration for males, yielding an MAE of 6.22 ± 0.58 years. However, this did not significantly outperform the second-best HGB male model in permutation tests (Data Aggregation multi-pipeline, multi-atlas concatenation; MAE = 6.27; p = 0.59).

Despite their higher dimensionality, cortical-augmented models remained less accurate than the base morphological models (best MAE 6.22 vs. 5.45). The key takeaway from this section is that neither atlas concatenation within a pipeline, nor pipeline aggregation across software versions yielded the gains observed when aggregating fundamentally distinct segmentation tools.

## 4. Discussion

### a) Analytical Variability Analyses

The analytical variability results reinforce that segmentation pipeline choice is not a neutral preprocessing decision. Inter-pipeline variability was spatially localized to structure boundaries rather than reflecting gross anatomical mislocalization. Mean voxelwise disagreement was extremely low, and fewer than 0.27% of voxels showed high disagreement (Supplementary Figure S1), indicating that pipelines broadly agree on sub-cortical anatomical cores. Discrepancies instead arise primarily from differences in boundary delineation.

Average DSCs varied substantially across pipeline pairs and structures, ranging from mediocre to very high. Across most comparisons, larger structures demonstrated higher spatial overlap, consistent with a size-related effect. However, this pattern was not consistent across all comparisons. In particular, attenuation of the correlation in the FSL–SAMSEG comparison suggests that algorithm-specific boundary conventions can counteract geometric size effects. While both pipelines show strong size-related trends when compared with FreeSurfer, their direct comparison reveals discrepancies that weaken the volume–overlap relationship. These findings indicate that apparent size effects in DSCs are partly geometric but also modulated by algorithmic differences.

Volume-based Spearman correlations and volumetric distribution analyses (Supplementary Figure S2) largely mirrored the DSC results: they were also highly variable across different pipelines, and smaller structures generally exhibited lower distributional agreement across pipelines. This pattern is consistent with surface-area–to–volume considerations, whereby small boundary shifts represent a larger proportion of total structure volume, degrading both spatial overlap and volumetric consistency. However, DSC and volumetric similarity were not interchangeable. Despite high mean DSC values, the thalamus demonstrated only moderate distributional overlap, illustrating how strong agreement in core voxels of large structures can obscure systematic boundary differences. This is particularly plausible at the thalamus–internal capsule interface, where low T1 contrast and subtle gray–white gradients may lead to differential treatment of partial-volume voxels. Together, these findings demonstrate that spatial overlap and volumetric similarity capture distinct dimensions of segmentation variability.

The RVD analysis further refines the interpretation of size effects. Although most pipeline pairings showed negative correlations between structure volume and proportional disagreement, only one remained significant after correction, and the overall association was modest. When disagreement is expressed relative to structure size, the apparent size dependence observed in DSC and raw volumetric comparisons largely attenuates. This suggests that smaller structures do not exhibit disproportionate proportional bias; instead, their lower DSC and volumetric agreement primarily reflects sensitivity of absolute metrics to small boundary shifts, rather than true scaling errors.

Overall, inter-pipeline variability appears driven more by pipeline-specific boundary definitions than by structure-size-dependent volumetric distortion, underscoring the importance of structure-level harmonization when integrating outputs across segmentation frameworks.

Importantly, inter-pipeline disagreement showed robust demographic associations. Age-related correlations ranged widely (r = −0.73 to 0.55), with both extremes observed in the thalamus. Negative correlations—where disagreement decreases with age—may indicate that certain pipeline atlas priors differ more strongly in younger anatomy but converge as age-related atrophy simplifies morphological variation. Conversely, positive correlations—frequently observed in the accumbens—suggest that aging amplifies methodological sensitivity, potentially due to increased atrophy, reduced contrast, or heightened boundary ambiguity. While most structures showed hemispheric symmetry, exceptions in the accumbens and putamen indicate lateralized interactions between aging and segmentation behavior.

Sex effects were also present, with larger disagreement more commonly observed in males. While RVDs were normalized by mean structure size, this adjustment may not entirely eliminate residual scaling effects associated with larger male head sizes. Consequently, these findings may reflect either subtle, uncorrected head-size influences or sex-specific interactions between morphology and segmentation algorithms. The largely symmetric hemispheric distribution of these effects further supports their systematic rather than stochastic nature.

Collectively, these findings demonstrate that segmentation pipeline selection meaningfully interacts with structure, age, and sex. Inter-pipeline variability is not random noise but exhibits systematic demographic sensitivity. Consequently, single-pipeline analyses risk conflating biological effects with algorithm-specific biases, particularly in small structures and age-heterogeneous samples.

Several limitations warrant consideration. The analysis was restricted to the 15 subcortical structures common across all pipelines, potentially underestimating variability in more granular parcellations such as hippocampal subfields or cortical structures. In the absence of manual gold-standard segmentations, inter-pipeline disagreement reflects methodological divergence rather than confirmed error. Technical differences in high-performance computing environments may introduce minor computational variability, although containerization minimized such effects. Finally, the female-skewed sample may limit generalizability of sex-specific disagreement patterns.

These results further underscore the importance of multi-pipeline validation when interpreting volumetric findings, particularly in small nuclei and demographically heterogeneous cohorts, where methodological variability may meaningfully intersect with biological inference.

### b) Brain Age Regression

The present findings demonstrate that feature concatenation across fundamentally distinct segmentation pipelines improves brain age prediction accuracy, whereas aggregation within or across versions of the same software framework does not yield comparable gains. In the base morphological models, combining outputs from different tools produced incremental but statistically significant improvements, particularly for tree-based learners. These gains likely reflect the intrinsic feature selection mechanisms of tree-based architectures, which are well suited to high-dimensional and partially redundant feature spaces. By prioritizing informative predictors and effectively down-weighting noisy or less reliable inputs, tree-based models can capitalize on multi-pipeline aggregation while mitigating the adverse effects of redundancy.

In contrast, linear (ElasticNet), distance-based (KNN), and kernel-based (SVM) models were less able to exploit aggregated feature sets. The inferior performance of the simple stacking ensemble further supports this interpretation. Because the ensemble relied on averaging predictions across pipelines, it lacked a mechanism to selectively weight higher-quality feature sources. The ensemble approach was likely compromised by incorporating systematically less reliable predictors without filtering or reweighting them. In this context, naive averaging appears insufficient to overcome pipeline-specific weaknesses.

Although the improvements achieved through concatenation were often modest in absolute terms, they were consistent and statistically significant. Moreover, concatenation generally enhanced stability across folds, suggesting that a more comprehensive and redundant multi-pipeline representation reduces sensitivity to fold-specific variability. This robustness likely arises from complementary signal capture across pipelines, where noise or bias in one processing stream is partially offset by others. This was more evident in male cohorts where baseline stability was lower.

In contrast, cortical-augmented models revealed diminishing returns from increasing feature dimensionality within a single software ecosystem. Neither atlas concatenation within FreeSurfer nor cross-version aggregation between FreeSurfer 7.4.1 and 8.0.0.1 produced consistent improvements. This suggests that predictive morphological signal across FreeSurfer versions is highly redundant. Although FreeSurfer 7 and 8 employ different segmentation algorithms, version 8 was partially trained on automated maps derived from earlier FreeSurfer releases, limiting algorithmic independence. Consequently, the substantial increase in dimensionality likely introduced noise that outweighed any incremental informational gains.

The superior performance of the base models relative to the cortical-augmented models further emphasizes the importance of algorithmic diversity over feature expansion. The base models combined fundamentally distinct segmentation tools—FreeSurfer, SAMSEG, and FSL—each trained on different datasets and modeling assumptions. This diversity appears more beneficial than simply increasing cortical granularity or combining closely related software versions. Additionally, the broad lifespan range of the cohort (6–90 years) may introduce morphological ambiguity in cortical metrics. Similar cortical surface area or thickness values may reflect either early-life developmental states or late-life atrophic processes, potentially reducing their discriminative utility for age prediction.

## 5. Conclusion

The findings of this study underscore the importance of multi-pipeline augmentation for brain age prediction. Leveraging a “multiverse” of features through multi-pipeline feature concatenation in our base models yielded the highest predictive accuracy, significantly outperforming any single-pipeline model. This result demonstrates that inter-pipeline variability is not merely methodological noise but can be harnessed as complementary signal. However, this benefit did not hold in cortical-augmented models, where neither within-version atlas concatenation nor cross-version pipeline aggregation consistently outperformed single-pipeline, single-atlas models.

These analyses also reveal the profound impact of analytical flexibility on subcortical volumetry. While individual pipelines demonstrate high spatial agreement in the anatomical core of structures, considerable variability at the boundaries drives volumetric discrepancies, particularly in smaller nuclei such as the accumbens and amygdala. These boundary-driven differences likely constitute a major source of downstream variation in predictive modeling and group-level inference.

Pipeline choice is not a demographically neutral decision. Inter-pipeline disagreement is significantly modulated by both age and sex, with males generally exhibiting higher disagreement and age-related atrophy often amplifying or masking methodological divergence. To mitigate the risk of analytical flexibility on the reproducibility crisis, we propose moving beyond single-pipeline analyses. We recommend that future neuroimaging studies check result robustness and bolster their findings by utilizing multiple processing pipelines. With the growing availability of deep learning–based segmentation models and the resulting reductions in processing time, incorporating multiple pipelines is becoming increasingly feasible even in large-scale studies. Furthermore, investigations should explicitly account for interactions between demographic covariates and segmentation behavior to provide a robust account of the biological factors at play.

## Supplementary Material

**Supplementary Table S1:** Hyperparameter Search Spaces for Machine Learning Models.

| Model | Hyperparameter | Search Space / Values Evaluated |
| --- | --- | --- |
| Elastic Net | Regularization Strength (alpha) | [0.001, 0.01, 0.1, 1.0, 10.0, 100.0] |
|  | L1 ratio | [0.1, 0.3, 0.5, 0.7, 0.9] |
| K-Nearest Neighbors | Number of neighbors (k) | [3, 5, 7, 9, 11] |
|  | Weight function | ['uniform', 'distance'] |
|  | Distance metric power (p) | [1, 2] |
| Support Vector Regression | Regularization parameter (C) | [0.01, 0.1, 1.0, 10.0, 100.0] |
|  | Kernel type | ['linear', 'rbf'] |
|  | Kernel coefficient (gamma) | ['scale', 'auto', 0.01, 0.1] |
| Extremely Randomized Trees | Number of trees | [100, 200] |
|  | Maximum depth | [10, 20, None] |
|  | Minimum samples to split | [2, 5] |
|  | Minimum samples at leaf | [1, 2] |
| Histogram-Gradient Boosting | Maximum iterations | [100, 200] |
|  | Learning rate | [0.01, 0.1] |
|  | Maximum leaf nodes | [31, 63] |
|  | Maximum depth | [5, None] |

**Figure S2:**
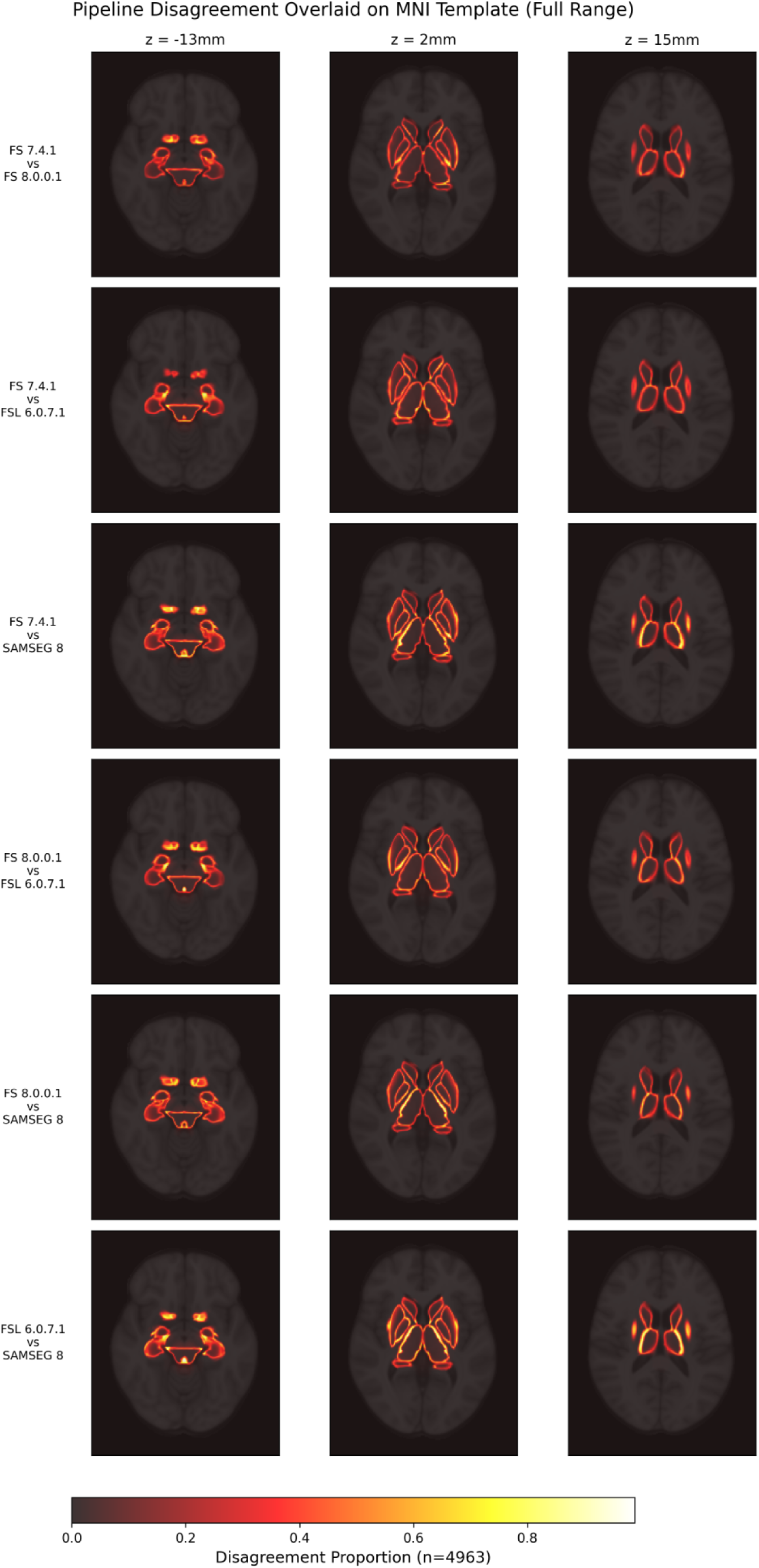
Heatmaps illustrating voxelwise subcortical segmentation disagreement across four pipelines, where colors represent the proportion of subjects with inter-pipeline variability across three representative axial slices.

Supplementary Figure S2 illustrates voxelwise disagreement across four pipelines using a multi-class comparison approach, capturing both structural boundary differences and internal label identity mismatches. Each voxel encodes the proportion of subjects (n=4963) for which pipelines assigned differing anatomical labels, displayed across three representative axial slices encompassing the 15 common subcortical structures. Across all pipeline pairs, agreement remained strong throughout the core volumes of the structures, with disagreement primarily localized at structural interfaces. Mean voxelwise disagreement was consistently low across comparisons (0.003-0.005), with a median of zero and fewer than 0.27% of voxels showing disagreement greater than 50%. High disagreement was consistently observed at both the outer boundaries of the subcortical mass and the internal interfaces between adjacent nuclei, particularly along the boundaries of the thalamus and basal ganglia. The spatial distribution and magnitude of these discrepancies were highly consistent across all evaluated pipeline pairs.

**Supplementary Figure S3:**
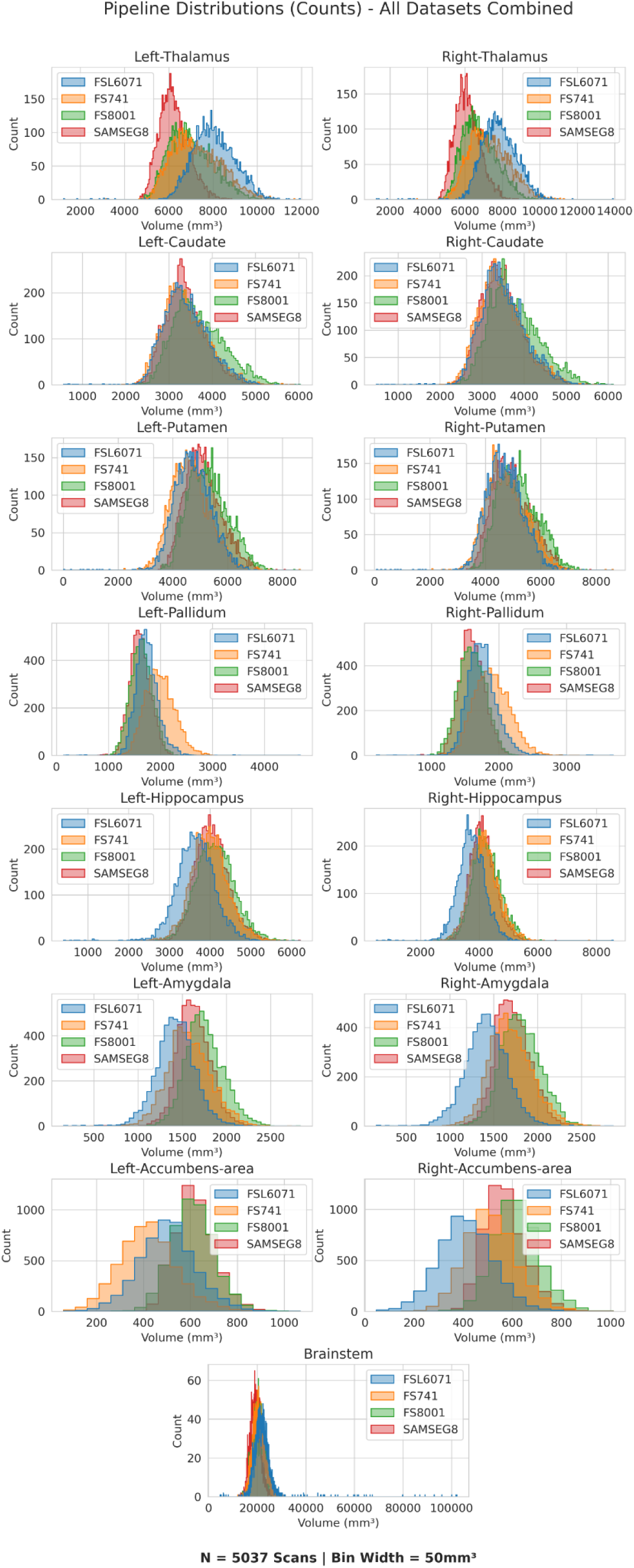
Volumetric distributions across the four segmentation pipelines for 15 subcortical structures.

Volumetric distributions of the 15 subcortical structures common across the four segmentation pipelines are shown in Figure 6. Inter-pipeline distribution similarity was assessed using two-sample Kolmogorov–Smirnov (K-S) tests for each structure across all pairwise pipeline comparisons (6 comparisons × 15 structures = 90 tests), with Bonferroni correction applied across all tests. For every structure, at least one pairwise comparison remained significant after correction (minimum adjusted p < 0.001), indicating that no structure exhibited complete distributional equivalence across pipelines.

To summarize agreement between pipelines, we report the mean 1−D value for each structure, where D is the K-S statistic (the maximum difference between empirical cumulative distribution functions). This measure represents the average distributional overlap across all 6 pairwise pipeline comparisons. Observed mean 1−D values ranged from 0.558 to 0.882. Consistent with the greater sensitivity of agreement metrics in smaller structures, larger structures—including the caudate (L/R: 0.876, 0.882), putamen (0.778, 0.845), hippocampus (0.776, 0.777), and brainstem (0.784)—showed relatively high overlap, whereas smaller structures such as the accumbens (0.566, 0.559), amygdala (0.672, 0.647), and pallidum (0.663, 0.698) had lower, though still moderate, agreement. However, the thalamus (0.601, 0.592) exhibited only moderate overlap despite its large anatomical size.

Systematic differences in central tendency were evaluated using two-sided Wilcoxon signed-rank tests for each pairwise pipeline comparison within each structure (90 tests total), with Bonferroni correction applied. A significant test indicates a systematic directional shift in volumes between the two pipelines for that structure. After correction, almost all comparisons remained significant: freesurfer8001 (45/45), fslanat6071 (44/45), freesurfer741 (44/45), and samseg8001 (45/45). The denominators (45) reflect the number of structure-specific pairwise comparisons involving each pipeline (15 structures × 3 comparisons per structure). Inspection of the significant pairwise comparisons reveals that no single pipeline consistently produced larger or smaller volumes across all structures. Notably, out of all 90 comparisons evaluated, only the right putamen volume comparison between FreeSurfer 7.4.1 and FSL Anat 6.0.1 failed to reject the null hypothesis after correction (adjusted p = 1.00).

## References

Beheshti, I., Maikusa, N., & Matsuda, H. (2022). The accuracy of T1-weighted voxel-wise and region-wise metrics for brain age estimation. Computer Methods and Programs in Biomedicine, 214, 106585. 10.1016/j.cmpb.2021.106585

Bhagwat, N., Barry, A., Dickie, E. W., Brown, S. T., Devenyi, G. A., Hatano, K., DuPre, E., Dagher, A., Chakravarty, M., Greenwood, C. M. T., Misic, B., Kennedy, D. N., & Poline, J.-B. (2021). Understanding the impact of preprocessing pipelines on neuroimaging cortical surface analyses. GigaScience, 10(1), giaa155. 10.1093/gigascience/giaa155

Bhagwat, N., Wang, M., Dugré, M., Pfarr, J.-K., Dai, A., Urchs, S., McPherson, B., Gau, R., Heese, E. M. van, d’Angremont, E., Laansma, M. A., Prasad, S., Sanz-Robinson, J., Torabi, M., Jahanpour, A., Danyluik, M., Joubert, A., Macdonald, A., Waller, L., … Poline, J.-B. (2026). Nipoppy: A framework for standardizing neuroimaging studies to facilitate international derived-data sharing (p. 2026.05.18.723593). bioRxiv. 10.64898/2026.05.18.723593

Botvinik-Nezer, R., Holzmeister, F., Camerer, C. F., Dreber, A., Huber, J., Johannesson, M., Kirchler, M., Iwanir, R., Mumford, J. A., Adcock, R. A., Avesani, P., Baczkowski, B. M., Bajracharya, A., Bakst, L., Ball, S., Barilari, M., Bault, N., Beaton, D., Beitner, J., … Schonberg, T. (2020). Variability in the analysis of a single neuroimaging dataset by many teams. Nature, 582(7810), Article 7810. 10.1038/s41586-020-2314-9

Botvinik-Nezer, R., & Wager, T. D. (2023). Reproducibility in Neuroimaging Analysis: Challenges and Solutions. *Biological Psychiatry: Cognitive Neuroscience and Neuroimaging*, Reliability of Neurocognitive Measures for Mental Health, 8(8), 780–788. 10.1016/j.bpsc.2022.12.006

Bowring, A., Maumet, C., & Nichols, T. E. (2019). Exploring the impact of analysis software on task fMRI results. Human Brain Mapping, 40(11), 3362–3384. 10.1002/hbm.24603

Capó, M., Vitali, S., Athanasiou, G., Cusimano, N., García, D., Cruickshank, G., & Patel, B. (2025). UK Biobank MRI data can power the development of generalizable brain clocks: A study of standard ML/DL methodologies and performance analysis on external databases. NeuroImage, 308, 121064. 10.1016/j.neuroimage.2025.121064

Carp, J. (2012). On the plurality of (methodological) worlds: Estimating the analytic flexibility of FMRI experiments. Frontiers in Neuroscience, 6, 149. 10.3389/fnins.2012.00149

CMA, M. G. H. (2003). General Brain Segmentation—Method and Utilization. The Center for Morphometric Analysis. https://www.nmr.mgh.harvard.edu/∼nikos/Public/CMA/CMA-Segmentation-Manual.pdf

Cole, J. H., Ritchie, S. J., Bastin, M. E., Valdés Hernández, M. C., Muñoz Maniega, S., Royle, N., Corley, J., Pattie, A., Harris, S. E., Zhang, Q., Wray, N. R., Redmond, P., Marioni, R. E., Starr, J. M., Cox, S. R., Wardlaw, J. M., Sharp, D. J., & Deary, I. J. (2018). Brain age predicts mortality. Molecular Psychiatry, 23(5), 1385–1392. 10.1038/mp.2017.62

De Bonis, M. L. N., Fasano, G., Lombardi, A., Ardito, C., Ferrara, A., Di Sciascio, E., & Di Noia, T. (2024). Explainable brain age prediction: A comparative evaluation of morphometric and deep learning pipelines. Brain Informatics, 11(1), 33. 10.1186/s40708-024-00244-9

Filipek, P. A., Richelme, C., Kennedy, D. N., & Caviness, V. S., Jr. (1994). The Young Adult Human Brain: An MRI-based Morphometric Analysis. Cerebral Cortex, 4(4), 344–360. 10.1093/cercor/4.4.344

Fischl, B. (2012). FreeSurfer. NeuroImage, 62(2), 774–781. 10.1016/j.neuroimage.2012.01.021

Fogarty, M. (2018). Subcortical Segmentation—Free Surfer Wiki. FreeSurfer.Net. https://freesurfer.net/fswiki/SubcorticalSegmentation

Franke, K., Ziegler, G., Klöppel, S., Gaser, C., & Alzheimer’s Disease Neuroimaging Initiative. (2010). Estimating the age of healthy subjects from T1-weighted MRI scans using kernel methods: Exploring the influence of various parameters. NeuroImage, 50(3), 883–892. 10.1016/j.neuroimage.2010.01.005

Glatard, T., Lewis, L. B., Ferreira da Silva, R., Adalat, R., Beck, N., Lepage, C., Rioux, P., Rousseau, M.-E., Sherif, T., Deelman, E., Khalili-Mahani, N., & Evans, A. C. (2015). Reproducibility of neuroimaging analyses across operating systems. Frontiers in Neuroinformatics, 9, 12. 10.3389/fninf.2015.00012

Gomez-Ramirez, J., Quilis-Sancho, J., & Fernandez-Blazquez, M. A. (2022). A Comparative Analysis of MRI Automated Segmentation of Subcortical Brain Volumes in a Large Dataset of Elderly Subjects. Neuroinformatics, 20(1), 63–72. 10.1007/s12021-021-09520-z

Guan, S., Jiang, R., Meng, C., & Biswal, B. (2024). Brain age prediction across the human lifespan using multimodal MRI data. GeroScience, 46(1), 1–20. 10.1007/s11357-023-00924-0

Kennedy, D. N., Abraham, S. A., Bates, J. F., Crowley, A., Ghosh, S., Gillespie, T., Goncalves, M., Grethe, J. S., Halchenko, Y. O., Hanke, M., Haselgrove, C., Hodge, S. M., Jarecka, D., Kaczmarzyk, J., Keator, D. B., Meyer, K., Martone, M. E., Padhy, S., Poline, J.-B., … Travers, M. (2019). Everything Matters: The ReproNim Perspective on Reproducible Neuroimaging. Frontiers in Neuroinformatics, 13. 10.3389/fninf.2019.00001

Kiar, G., Chatelain, Y., Salari, A., Evans, A. C., & Glatard, T. (2021). *Data Augmentation Through Monte Carlo Arithmetic Leads to More Generalizable Classification in Connectomics* (arXiv:2109.09649). arXiv. 10.48550/arXiv.2109.09649

Kiar, G., Mumford, J. A., Xu, T., Vogelstein, J. T., Glatard, T., & Milham, M. P. (2024). Why experimental variation in neuroimaging should be embraced. Nature Communications, 15, 9411. 10.1038/s41467-024-53743-y

Korbmacher, M., Westlye, L. T., & Maximov, I. I. (2024). FreeSurfer version-shuffling can enhance brain age predictions. NeuroImage: Reports, 4(3), 100214. 10.1016/j.ynirp.2024.100214

Lao, Z., Shen, D., Xue, Z., Karacali, B., Resnick, S. M., & Davatzikos, C. (2004). Morphological classification of brains via high-dimensional shape transformations and machine learning methods. NeuroImage, 21(1), 46–57. 10.1016/j.neuroimage.2003.09.027

Leys, C., Ley, C., Klein, O., Bernard, P., & Licata, L. (2013). Detecting outliers: Do not use standard deviation around the mean, use absolute deviation around the median. Journal of Experimental Social Psychology, 49(4), 764–766. 10.1016/j.jesp.2013.03.013

Litwińczuk, M. C., Muhlert, N., Trujillo-Barreto, N., & Woollams, A. (2024). Impact of brain parcellation on prediction performance in models of cognition and demographics. Human Brain Mapping, 45(2), e26592. 10.1002/hbm.26592

Liu, S., Xu, L., Xu, G., Wang, Y., Zhang, G., & He, L. (2025). A systematic review: Brain age gap as a promising early diagnostic biomarker for Alzheimer’s disease. Journal of the Neurological Sciences, 475, 123563. 10.1016/j.jns.2025.123563

Modabbernia, A., Whalley, H. C., Glahn, D. C., Thompson, P. M., Kahn, R. S., & Frangou, S. (2022). Systematic evaluation of machine learning algorithms for neuroanatomically-based age prediction in youth. Human Brain Mapping, 43(17), 5126–5140. 10.1002/hbm.26010

Nastase, S. A., Liu, Y.-F., Hillman, H., Zadbood, A., Hasenfratz, L., Keshavarzian, N., Chen, J., Honey, C. J., Yeshurun, Y., Regev, M., Nguyen, M., Chang, C. H. C., Baldassano, C., Lositsky, O., Simony, E., Chow, M. A., Leong, Y. C., Brooks, P. P., Micciche, E., … Hasson, U. (2021). The “Narratives” fMRI dataset for evaluating models of naturalistic language comprehension. Scientific Data, 8(1), 250. 10.1038/s41597-021-01033-3

Nooner, K. B., Colcombe, S., Tobe, R., Mennes, M., Benedict, M., Moreno, A., Panek, L., Brown, S., Zavitz, S., Li, Q., Sikka, S., Gutman, D., Bangaru, S., Schlachter, R. T., Kamiel, S., Anwar, A., Hinz, C., Kaplan, M., Rachlin, A., … Milham, M. (2012). The NKI-Rockland Sample: A Model for Accelerating the Pace of Discovery Science in Psychiatry. Frontiers in Neuroscience, 6. 10.3389/fnins.2012.00152

Nugent, A. C., Thomas, A. G., Mahoney, M., Gibbons, A., Smith, J. T., Charles, A. J., Shaw, J. S., Stout, J. D., Namyst, A. M., Basavaraj, A., Earl, E., Riddle, T., Snow, J., Japee, S., Pavletic, A. J., Sinclair, S., Roopchansingh, V., Bandettini, P. A., & Chung, J. (2022). The NIMH intramural healthy volunteer dataset: A comprehensive MEG, MRI, and behavioral resource. Scientific Data, 9(1), 518. 10.1038/s41597-022-01623-9

Patenaude, B., Smith, S. M., Kennedy, D. N., & Jenkinson, M. (2011). A Bayesian model of shape and appearance for subcortical brain segmentation. NeuroImage, 56(3), 907–922. 10.1016/j.neuroimage.2011.02.046

Puonti, O., Iglesias, J. E., & Van Leemput, K. (2016). Fast and sequence-adaptive whole-brain segmentation using parametric Bayesian modeling. NeuroImage, 143, 235–249. 10.1016/j.neuroimage.2016.09.011

Rajabli, R., Soltaninejad, M., Fonov, V. S., Bzdok, D., & Collins, D. L. (2025). Brain Age Prediction: Deep Models Need a Hand to Generalize. Human Brain Mapping, 46(11), e70254. 10.1002/hbm.70254

Sanz-Robinson, J., Jahanpour, A., Phillips, N., Glatard, T., & Poline, J.-B. (2022). NeuroCI: Continuous Integration of Neuroimaging Results Across Software Pipelines and Datasets. 2022 IEEE 18th International Conference on E-Science (e-Science), 105–116. 10.1109/eScience55777.2022.00025

Schilling, K. G., Rheault, F., Petit, L., Hansen, C. B., Nath, V., Yeh, F.-C., Girard, G., Barakovic, M., Rafael-Patino, J., Yu, T., Fischi-Gomez, E., Pizzolato, M., Ocampo-Pineda, M., Schiavi, S., Canales-Rodríguez, E. J., Daducci, A., Granziera, C., Innocenti, G., Thiran, J.-P., … Descoteaux, M. (2021). Tractography dissection variability: What happens when 42 groups dissect 14 white matter bundles on the same dataset? (p. 2020.10.07.321083). 10.1101/2020.10.07.321083

Sokołowski, A., Bhagwat, N., Kirbizakis, D., Chatelain, Y., Dugré, M., Poline, J.-B., Sharp, M., & Glatard, T. (2024). The impact of FreeSurfer versions on structural neuroimaging analyses of Parkinson’s disease (p. 2024.11.11.623071). bioRxiv. 10.1101/2024.11.11.623071

Spreng, R. N., Setton, R., Alter, U., Cassidy, B. N., Darboh, B., DuPre, E., Kantarovich, K., Lockrow, A. W., Mwilambwe-Tshilobo, L., Luh, W.-M., Kundu, P., & Turner, G. R. (2022). Neurocognitive aging data release with behavioral, structural and multi-echo functional MRI measures. Scientific Data, 9(1), 119. 10.1038/s41597-022-01231-7

Villeneuve, S., Poirier, J., Breitner, J. C. S., Tremblay-Mercier, J., Remz, J., Raoult, J.-M., Yakoub, Y., Gallego-Rudolf, J., Qiu, T., Fajardo Valdez, A., Mohammediyan, B., Javanray, M., Metz, A., Sanami, S., Ourry, V., Wearn, A., Pastor-Bernier, A., Edde, M., Gonneaud, J., … Group, the P.-A. R. (2025). The PREVENT-AD cohort: Accelerating Alzheimer’s disease research and treatment in Canada and beyond. Alzheimer’s & Dementia, 21(10), e70653. 10.1002/alz.70653

Wu, Y., Gao, H., Zhang, C., Ma, X., Zhu, X., Wu, S., & Lin, L. (2024). Machine Learning and Deep Learning Approaches in Lifespan Brain Age Prediction: A Comprehensive Review. Tomography, 10(8), 1238–1262. 10.3390/tomography10080093

Yücel, M. A., Luke, R., Mesquita, R. C., von Lühmann, A., Mehler, D. M. A., Lührs, M., Gemignani, J., Abdalmalak, A., Albrecht, F., de Almeida Ivo, I., Artemenko, C., Ashton, K., Augustynowicz, P., Bajracharya, A., Bannier, E., Barth, B., Bayet, L., Behrendt, J., Khani, H. B., … Zemanek, V. (2024). *The fNIRS Reproducibility Study Hub (FRESH): Exploring Variability and Enhancing Transparency in fNIRS Neuroimaging Research* (Pc6x8_v1). MetaArXiv. 10.31222/osf.io/pc6x8_v1

Zeighami, Y., & Evans, A. C. (2021). Association vs. Prediction: The Impact of Cortical Surface Smoothing and Parcellation on Brain Age. Frontiers in Big Data, 4. 10.3389/fdata.2021.637724

Zhang, R., Yi, F., Mao, H., Huang, Z., Wang, K., & Zhang, J. (2025). Brain age gap as a predictive biomarker that links aging, lifestyle, and neuropsychiatric health. Communications Medicine, 5(1), 441. 10.1038/s43856-025-01100-5

Zhao, T., Liao, X., Fonov, V. S., Wang, Q., Men, W., Wang, Y., Qin, S., Tan, S., Gao, J.-H., Evans, A., Tao, S., Dong, Q., & He, Y. (2019). Unbiased age-specific structural brain atlases for Chinese pediatric population. NeuroImage, 189, 55–70. 10.1016/j.neuroimage.2019.01.006

